# Fast exploration is coupled with a less choosy but more reactive learning style in a generalist predator

**DOI:** 10.1101/2024.03.26.586890

**Authors:** Chi-Yun Kuo, Yu-Hsi Chen, Ai-Ching Meng, Yu-Zhe Wu, Shan-Yu Yang, Ching-Ning Yeh

## Abstract

The hypothesis of slow-fast syndromes predicts a correspondence between personality type and learning style; fast explorers would have a more proactive (fast but inflexible) learning style and slow explorers would be more reactive (slow but flexible) learners. Empirical evidence for this personality-cognition coupling remains inconclusive and heavily biased towards birds. Moreover, most studies did not examine the personality-cognition correlation when the cognitive task is discerning food quality, a scenario directly related to energy acquisition that underpins the evolution of slow-fast syndromes. In this study, we examined the exploration-cognition correlation in the context of avoidance learning in an opportunistic predator - the common sun skink *Eutropis multifasciata*. We quantified exploration tendencies of individuals in an unfamiliar environment and compared foraging behaviours when lizards associated prey colour and quality during the initial learning trials and subsequent reverse learning trials, where the prey colour-taste combinations were switched. We found that fast explorers were less choosy but more reactive foragers, whereas slow explorers exhibited the opposite learning style. Interestingly, there was no evidence for a learning speed-flexibility trade-off. Our findings are in contrast with conventional predictions and suggest that the two types of exploration-cognition coupling could be different viable responses to fast-changing environmental predictability.

## Introduction

Consistent behavioural tendencies within and across contexts (i.e., animal personalities or temperaments) have been widely recognized as an important mechanism for maintaining individual-level behavioural variation [1,2]. Over the past two decades, the expression of personalities has been shown to covary with other traits, such as nutritional state and metabolic rate [3,4], and examined in larger contexts such as life-history theory [5]. One major motivation of examining the covariation between personality and other organismal traits is to reveal trade-offs that underlie the formation of the fast-slow “syndrome” [6]. In short, fast individuals are more risk-prone (e.g., bolder, more explorative, less choosy, etc) and prioritize on short-term gains, whereas slow individuals tend to exhibit the opposite trait combinations and priorities. Along this vein, a notable hypothesis is the adaptive correlation between personality and cognition, which argues that selection would produce a similar and correlated continuum of cognitive styles along with personalities [7,8]. The idea behind the personality-cognition coupling is that the fast-slow continuum in behavioural types would covary with variation in cognitive styles mediated by an overarching risk-reward trade-off. Under this hypothesis, fast individuals, due to their risk-prone tendencies, would adopt or evolve a cognitive style characterized by faster, less choosy, less accurate, and less flexible learning, and the opposite would be true for slow individuals. Such associations are hypothesized to result from cognitive constraints and would benefit fast individuals in more stable environments and give slow individuals an advantage in more unpredictable environments [7].

The existence of personality-cognition coupling as predicted by a fast-slow syndrome has been tested in a substantial number of studies (reviewed in [9]). However, the empirical results overall did not support the aforementioned pattern, suggesting that personality-cognition correlations along the fast-slow continuum might be more complex than previously conceptualized. Notably, there are two major knowledge gaps from the literature. First, available studies are heavily biased towards birds, whereas taxa such as non-avian reptiles and insects have received far less attention. This bias in taxonomic representation can have important consequences, as these groups may modify behaviour with experience in ways that are more nuanced than birds. For example, a previous study showed that, unlike birds, foraging sequence rather than overall avoidance was the main foraging behaviour that a generalist lizard predator modified when associating prey colour and quality during avoidance learning trials [10]. Second, the personality-cognition coupling has rarely been examined in situations where animals must learn to discern food quality (e.g., [11,12]), especially when avoidance learning is the cognitive task (e.g., [13]), even though these scenarios directly relate to energy acquisition, which underpins the evolution and maintenance of a fast-slow syndrome [4,5]. We therefore posit that the personality-cognition correlation may be more detectable in contexts that involve foraging decision-making.

This study aims to fill the abovementioned knowledge gaps by examining the correlation between exploration tendencies and avoidance learning abilities in a widely foraging, generalist lizard predator. It has been proposed that widely foraging species, due to their higher activity level, tend to be exposed to higher mortality risks, experience higher rates of energy turnover, and encounter a wider range of prey items (e.g., [14–16]). These particular sets of conditions might be more conducive to a tighter coupling between exploration-related personalities and foraging-related cognitive abilities, compared to more sit-and-way species. We focus on two aspects of foraging decision-making that can be modified by learning – overall level of avoidance towards low-quality food items and foraging priority. Overall level of avoidance, commonly quantified as the proportion of unprofitable food items attacked or consumed, is the foraging outcome measured in the vast majority of avoidance learning experiments (e.g., [17–20]). Foraging priority, which describes the tendency to attack or consume more profitable food items first during a foraging sequence, regardless of the overall level of avoidance, is a largely overlooked yet important aspect of foraging behaviour [10]. The focal cognitive traits are learning speed and learning flexibility. In the context of this study, the former describes the speed at which individuals adaptively change their foraging behaviour when first associating prey colour with prey quality, and the latter describes learning performance during a subsequent “reversal” phase when the prey colour-quality combination is switched [9].

Specifically, we test the following predictions. First, we predict that cognitive constraints will impose a trade-off between learning speed and learning flexibility, such that individuals cannot maximize both simultaneously, irrespective of their exploration tendencies. Second, more exploratory individuals (i.e., fast explorers) will be less choosy towards low quality prey while being faster yet more inflexible learners, whereas slow explorers will possess the opposite cognitive characteristics. Data in line with the predictions would support the existence of personality-cognition syndrome as hypothesized by [7]. Alternatively, if the exploration-learning correlation is partially supported (e.g., only learning speed or learning flexibility is associated with exploration in the predicted manner), it would suggest that learning speed and flexibility might not be constrained by cognitive trade-offs and could respond independently to the environment. If the correlation between exploration and learning is in the direction opposite to the prediction (e.g., fast explorers are slow but more flexible learners), it would suggest that selection might favour a different coupling between exploration and avoidance learning in foraging contexts.

## Materials and Methods

### Study species and husbandry

Our focal species is the common sun skink *Eutropis multifasciata*. This species is native to south and south-eastern Asian tropics but has expanded their geographic distributions in recent decades due to natural or human-assisted invasions. *Eutropis multifasciata* is a widely foraging predator with one of the most diverse diets among tropical skinks [21,22]. They are opportunistic predators whose diet can vary substantially seasonally and between regions [21,22]. We collected lizards from a population in Southern Taiwan (22.48ºN, 120.57ºE) from September to December of 2023. We determined the sexual maturity and the sex of the individuals based on external morphology and published adult snout-to-vent lengths (SVL) of 99 mm for males and 96 mm for females [23]. Individuals were brought back to the lab on the same day of capture and housed individually (cage dimension: 30 × 20 × 25 cm) under a 12L:12D cycle and temperatures of 28-30ºC. Each cage had beechwood bedding, a ceramic block for thermoregulation and a shelter. We provided the lizards with fresh water and food (crickets or meal worms) daily except during avoidance learning experiments (see below). We collected data from a total of 23 lizards (12 female and 11 males).

### Exploration assays

We quantified exploration tendencies of *E. multifasciata* with a novel environment design. We assayed each individual on the first and again on the seventh day after capture to confirm the repeatability of exploration. Assays were performed in a terrarium (length × width × height: 90 × 30 × 45 cm) with beechwood chips bedding. An opaque division separated the terrarium into two areas (30 or 60 × 30 × 45 cm). Prior to the assay, we painted a white dot on the lizard’s back with water-insoluble paint and transferred it from its home cage into the smaller chamber to acclimate for 10 mins. The smaller area contained a shelter and the ceramic block from the lizard’s home cage, representing a more familiar and safer space, whereas the larger, empty area represents the novel environment which the lizard could explore. We removed the division after the acclimation period and recorded the lizard’s behaviour for 30 mins with a GoPro camera mounted one meter above the terrarium (Hero 8, GoPro, San Matteo, CA, USA). We sprayed the shelter, the bedding, and the terrarium walls with 70% ethanol after each trial to minimize the potential influence of residual chemicals on the next lizard tested.

From each video, we manually extracted the following three variables: moving latency, exploration latency, and percent time in the novel environment. Moving latency was the time in seconds between the removal of the division to the time when the lizard made the first directional movement. Exploration latency was the time in seconds before at least half of the lizard’s body moved into the larger area. If this never happened for a lizard, we assigned a value of 2000 to this variable. Percent time in the novel environment was calculated as time in seconds during which at least half of the lizard’s body was in the novel environment divided by total observation time. In addition, we trained the software DeepLabCut [24] to automatically identify the location of the white dot on the lizard from each video frame (x- and y-coordinates in unit of pixel). We then calculate the total travel distance of the lizard during the assay based on the location of the white dot and divided the value by total observation time for standardization. This value, standardized distance travelled, is the fourth variable we used to quantify exploration.

### Avoidance learning experiments

Avoidance learning experiments had a learning phase and a reverse-learning phase. The learning phase consisted of five daily learning trials, in which lizards were presented with crickets that differed consistently in colour (red or yellow) and taste (normal or bitter), resulting in two colour-taste combination treatments (red-bitter and yellow-normal vs. red-normal and yellow-bitter). These learning trials allowed the lizards to associate prey colour with taste and modified their foraging behaviours accordingly. The learning trails were immediately followed by five daily reverse-learning trials in which the colour-taste combinations were switched. Seven females and six males encountered the former colour-taste combination during the learning phase, and five females and five males encountered the latter. Lizards were fasted for two days before the onset of avoidance learning experiments. We manipulated the colour of the crickets by painting their abdomens using water insoluble acrylic paint (MONA Acrylic Colour S-209 for the yellow colour and S-203 for the red colour). Bitter crickets were injected with ∼ 0.3 ml of 0.07% aqueous denatonium benzoate solution into the crickets, and we injected the normal-tasting crickets with a similar amount of water. This bitterness treatment was effective in eliciting strong negative reactions and avoidance learning in this species [10].

In each trial, we recorded the interactions between the focal lizard and the crickets for 15 minutes or until the lizard had attacked all crickets. An interaction between the lizard and the cricket was recorded as an “attack” if there was an unambiguous foraging attempt from the lizard. If the cricket was subsequently swallowed by the lizard, we recorded the interaction as “consume”. A “consume” was therefore always preceded by an “attack”. From the interactions, we calculated two variables that represented two important aspects of foraging decision-making – foraging priority and level of avoidance [10]. Foraging priority was calculated as the percentage of bitter crickets among the first 50% crickets attacked. A value of zero meant that a lizard attacked all normal crickets before attacking the bitter crickets, and a value of one denoted the opposite priority. We only calculated this variable if the focal lizard attacked at least four crickets in a trial. We calculated the level of avoidance as the percentage of bitter crickets among all crickets consumed in a trial. A value close to 0.5 would indicate that the lizard consumes normal and bitter crickets indiscriminately. We removed all remaining crickets at the end of each trial.

### Statistical analyses

We performed all statistical analyses in R (version 4.3.1, R Core Team).

To summarize individuals’ exploration tendencies, we used a principal component analysis (PCA) on moving latency, exploration latency, percent time in the novel environment, and standardized distance travelled to reduce the dimension of data. To test the repeatability of exploration tendency between trials, we built three generalized linear mixed models with individual identity as the random factor. The first model did not include any fixed factors. The second and third models included sex and trial as fixed factors, but only the second model included an interaction term. We used the Akaike Information Criterion (AIC) to select the optimal model and followed the model validation procedure recommended by [25] before interpretation.

To test the correlation between exploration and learning abilities, we constructed separate generalized linear models (GLM) with foraging priority or level of avoidance as response variables, using data from all trials. Grubb’s test revealed no potential outliers. This resulted in a sample size of 183 for foraging priority, after excluding trials in which the lizard attacked fewer than four crickets, and a sample size of 213 for level of avoidance, excluding trials in which the lizard did not consume any cricket. Since foraging priority and level of avoidance were proportions derived from count data, we used the binomial distribution with the complementary log-log link function in our GLMs. We included sex, colour-taste treatment, and trial as fixed factors. Individual identity was not included in the statistical models as the random effect variance was trivial (< 0.0001). Visual examination of data showed potential interactions between factors, so we used a model selection approach based on AIC to find the optimal model for interpretation. We started with a model with all possible two-way interactions between fixed effect factors and allowed only backward elimination, using the *step* function in the R package *stats*. We did not consider three-way interactions for its difficulty of interpretation and the small number of observations under each three-way intersection of data. We validated each optimal model based on [25] using the *DHARMa* package [26]. We presented all results regarding p-values using the language of evidence instead of a significance dichotomy [27]. In addition, we reported Cohen’s D for individual factors using the *effectsize* package [28] and interpreted the magnitude of the effects based on [29].

## Results

The first principal component (PC) summarized 65% of the total variation and we used the PC score to represent individuals’ exploration tendencies (table 1). Individuals with more negative PC scores moved more, spent more time in the novel environment, took less time to make the first directional movement, and were sooner to enter the novel environment. The difference in AIC scores between all three models was trivial (185.33 vs. 185.52 vs. 185.31), so we chose the full model with sex, trial, and the interaction term as fixed effect factors for validation and interpretation. Based on the results, there was no evidence that exploration tendencies differed between sexes and the two trials (table 2). We therefore used the average PC scores from the two trials to represent an individual’s overall exploration tendency in subsequent analyses.

**Table 1.** Factor loadings of the first principal component summarizing the variance in exploratory behaviours in the common sun skinks *Eutropis multifasciata*.

| Behavioural variables | PC1 |
| --- | --- |
| Moving latency | 0.505 |
| Exploration latency | 0.583 |
| Standardized distance travelled | -0.369 |
| Percent time in the novel environment | -0.518 |
| Cumulative % variance | 0.653 |

**Table 2.**
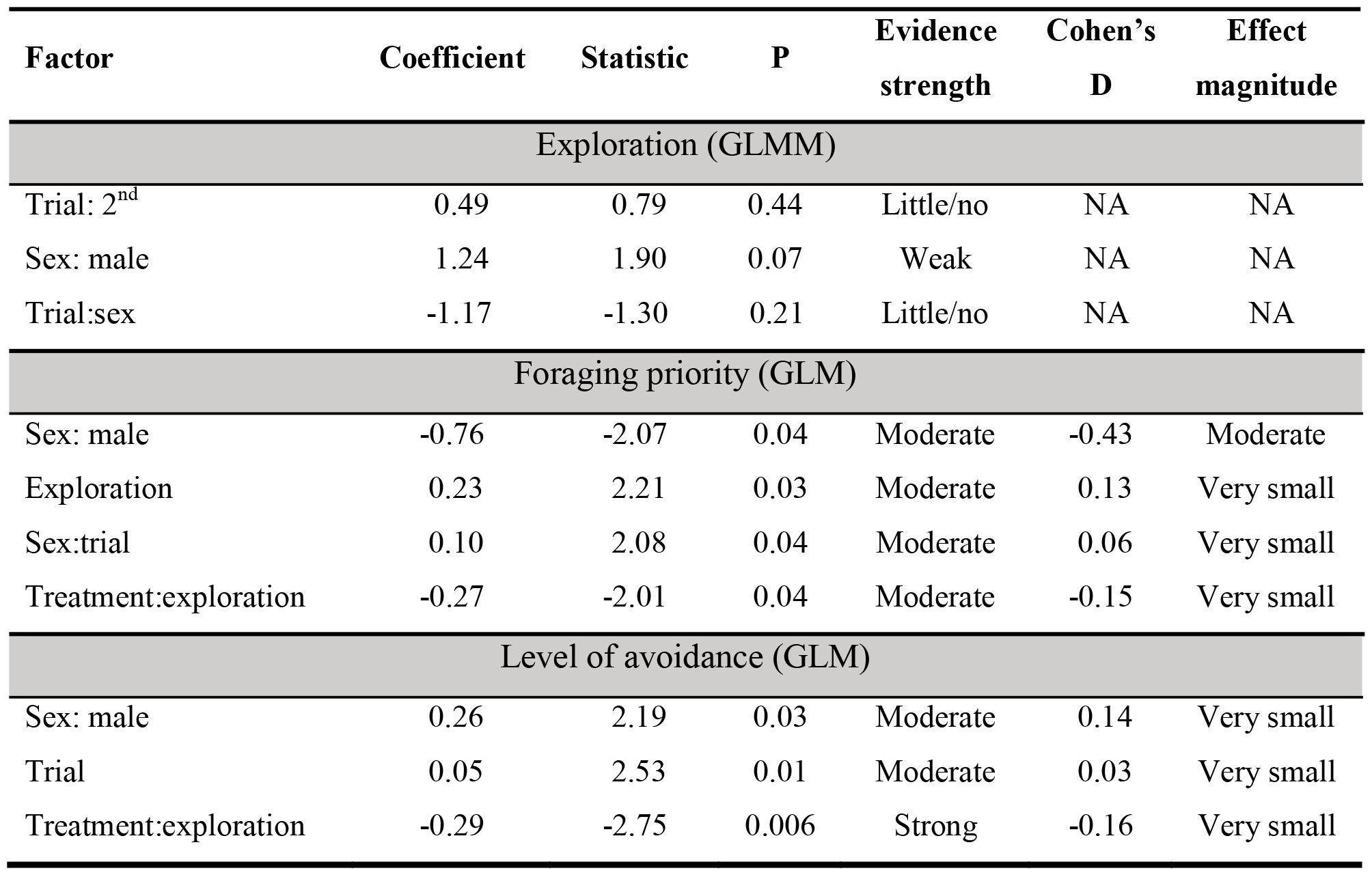
Summary of statistical analyses. Results were from GLMM for the analysis on the repeatability of and variation in exploration and GLM for analyses on the personality-cognition coupling. Due to space limitation, only factors with P < 0.05 were presented from GLM. It was t-statistic for the GLMM and z-statistic for the GLM. Interpretations of evidence strength were based on [27] and effect size magnitudes were based on [29]. YR treatment meant bitterness was first associated with yellow (Y) in the leaning phase and then red (R) in the reverse learning phase.

| Factor | Coefficient | Statistic | P | Evidence strength | Cohen's D | Effect magnitude |
| --- | --- | --- | --- | --- | --- | --- |
| Exploration (GLMM) |  |  |  |  |  |  |
| Trial: 2<sup>nd</sup> | 0.49 | 0.79 | 0.44 | Little/no | NA | NA |
| Sex: male | 1.24 | 1.90 | 0.07 | Weak | NA | NA |
| Trial:sex | -1.17 | -1.30 | 0.21 | Little/no | NA | NA |
| Foraging priority (GLM) |  |  |  |  |  |  |
| Sex: male | -0.76 | -2.07 | 0.04 | Moderate | -0.43 | Moderate |
| Exploration | 0.23 | 2.21 | 0.03 | Moderate | 0.13 | Very small |
| Sex:trial | 0.10 | 2.08 | 0.04 | Moderate | 0.06 | Very small |
| Treatment:exploration | -0.27 | -2.01 | 0.04 | Moderate | -0.15 | Very small |
| Level of avoidance (GLM) |  |  |  |  |  |  |
| Sex: male | 0.26 | 2.19 | 0.03 | Moderate | 0.14 | Very small |
| Trial | 0.05 | 2.53 | 0.01 | Moderate | 0.03 | Very small |
| Treatment:exploration | -0.29 | -2.75 | 0.006 | Strong | -0.16 | Very small |

For foraging priority, there was strong evidence for a very small effect of exploration and an interaction between exploration and colour treatment (table 2). The shift in foraging priority between the fifth and sixth trials indicated that lizards were confused by the sudden switch in the colour-taste association (figure 1). Regardless of colour treatment, slow and fast explorers modified their foraging priorities in contrasting ways in the learning phase (figure 1). Even though all individuals quickly learned to prioritize on normal crickets in the first trial, fast explorers gradually reversed this priority, no longer prioritizing on normal crickets at the end of the learning phase (figure 2b, d). Slow explorers, on the other hand, maintained this priority throughout the learning phase (figure 2a, c). All individuals learned to prioritize on normal crickets in the reverse learning phase, with the exception of slow explorers that encountered yellow-bitter combination (figure 1). These individuals still prioritized on yellow crickets despite their bitter taste (figure 1a). In addition, we also saw moderate evidence that sex played a mediating role in the modification of foraging priorities, with females being choosier and prioritizing on normal crickets more in the reverse learning phase (table 2, figure 3a, b).

**Figure 1.**
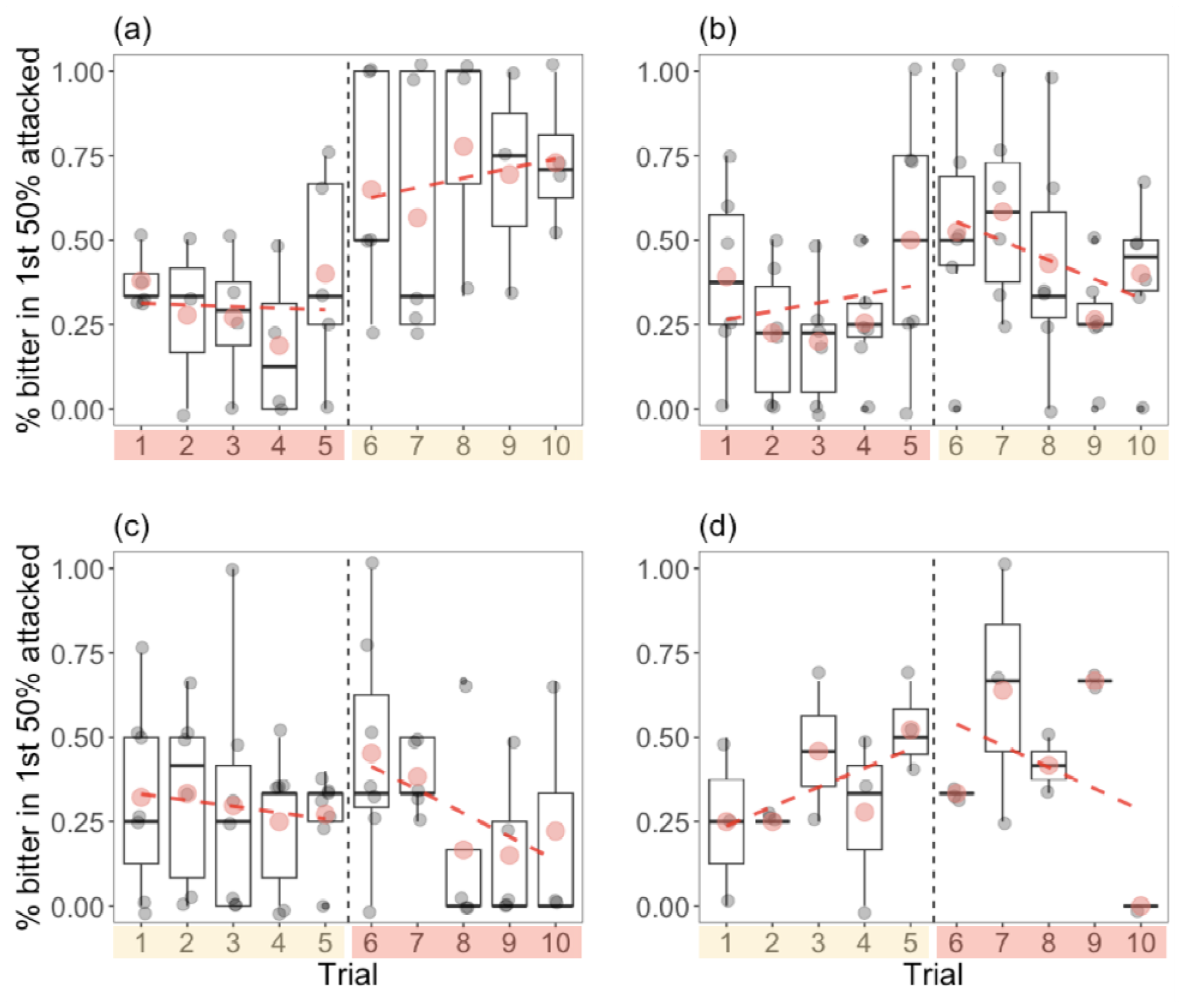
The correlation between exploration tendency and changes in foraging priority during the learning (trials one to five) and reverse learning (trials six to ten) in the common sun skinks, mediated by colour treatment. Gray circles are data from individuals, red circles are means within trials, and red dotted lines are linear regressions based on trial means to better illustrate the general trend. We defined slow explorers as having lower exploration PC scores than the population median and fast explorers as the opposite. Note that this categorization is intended to aid visual presentation of data; we treated exploration as a continuous variable in the formal analyses. Shaded boxes denote the colour associated with bitterness in these trials. (a) and (c) are data from slow explorers, and (b) and (d) are data from fast explorers.

**Figure 2.**
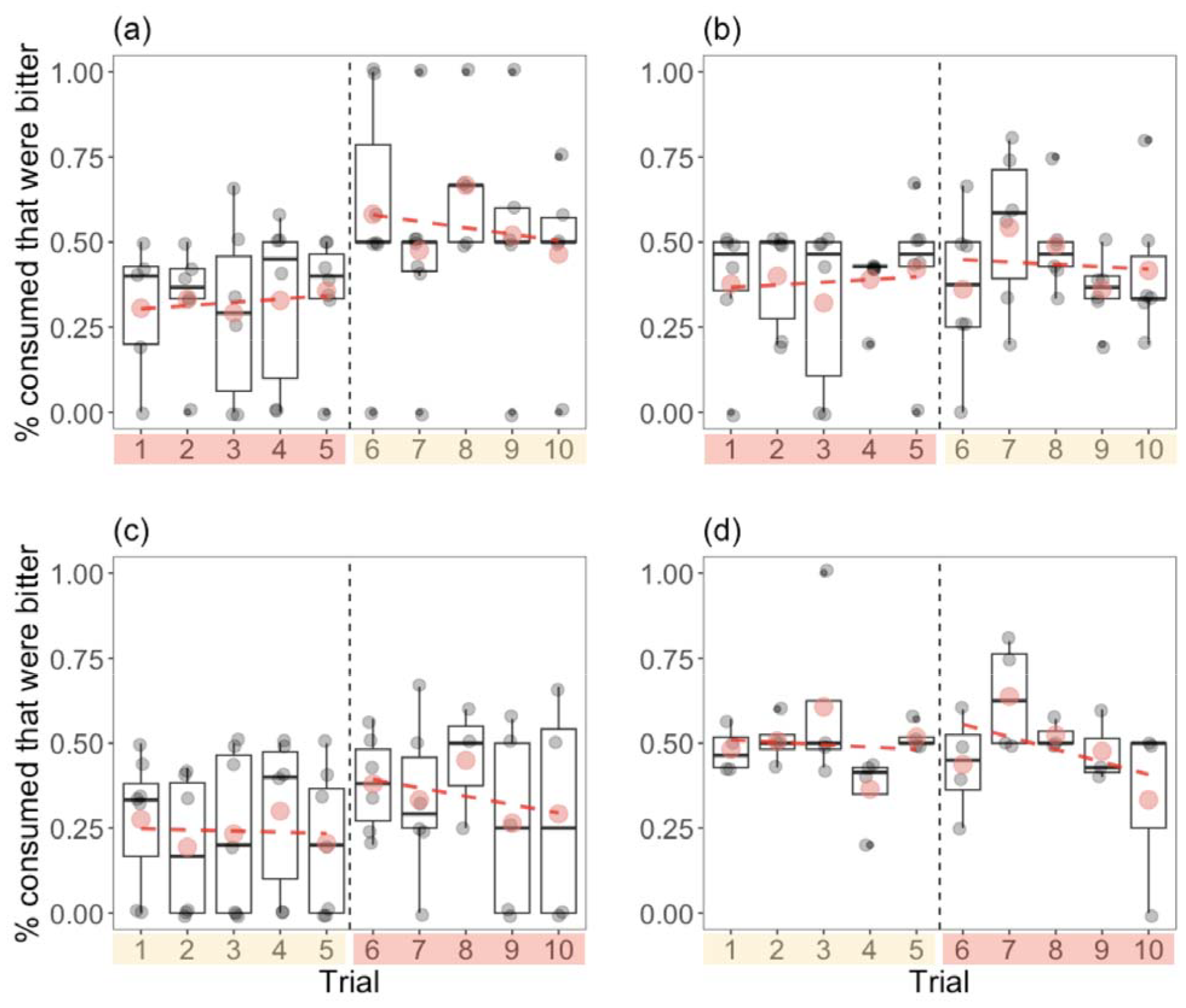
The correlation between exploration tendency and changes in level of avoidance during the learning (trials one to five) and reverse learning (trials six to ten) in the common sun skinks, also mediated by colour treatment. (a) and (c) are data from slow explorers, and those in (b) and (d) are from fast explorers. Coloured circles, dotted lines, and shaded boxes convey the same information as in figure 1.

**Figure 3.**
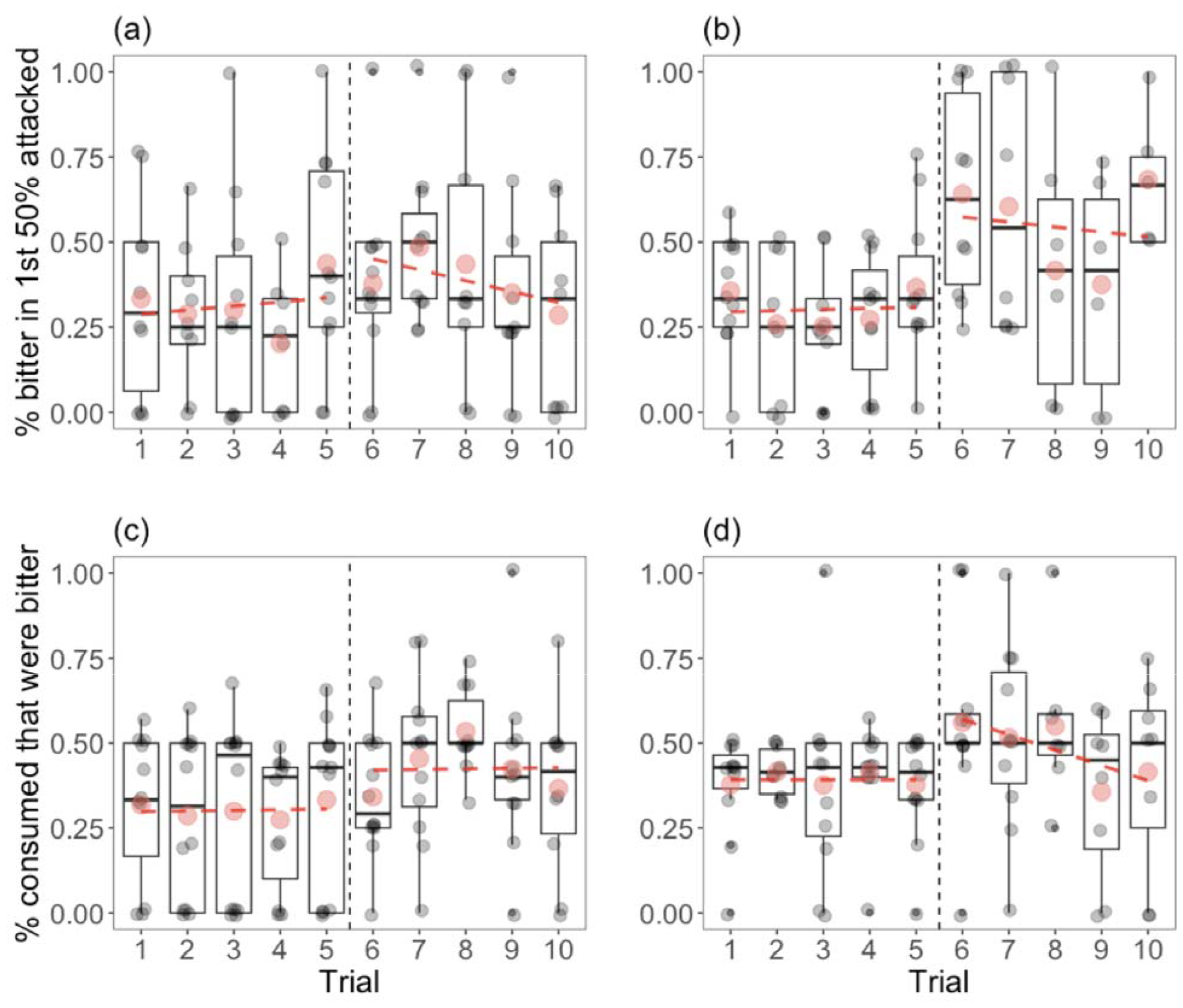
Sexual differences in how foraging behaviours are modified with learning. (a) and (b) are data for foraging priority, and (c) and (d) are data for level of avoidance. (a) and (c) show data from females, and (b) and (d) show those from males. Coloured circles and dotted lines convey the same information as in figures 1 and 2.

Regarding level of avoidance, there was strong evidence for a very small interactive effect of exploration and colour treatment (table 2). Similar to foraging priority, there was a decrease in avoidance between the fifth and the sixth trials, indicating the effectiveness of our experimental design (figure 2). Slow explorers by and large showed higher avoidance towards bitter crickets, regardless of experimental phase and colour treatment (figure 2a, c). The notable exception was slow explorers that encountered red- and then yellow-bitter combinations; these individuals still avoided red crickets to some extent in the reverse learning trials, leading to a lower avoidance towards yellow, bitter crickets (figure 2a). By contrast, fast explorers consumed normal and bitter crickets more indiscriminately, especially those that associated first yellow and then red with bitterness (figure 2b, d). Due to their lower choosiness, faster explorers consumed more crickets per trial throughout the experiment (5.93 vs. 4.38, one-way ANOVA, F_1,211_ = 35.93, p < 0.0001).

Even though the evidence strengths of our findings were at least moderate, we acknowledge that the effect magnitudes were small. Given the high variability typically associated with raw learning data (e.g., [17,30,31]), we argue that the small effect sizes were more the consequence of the inherent noise in the data rather than the triviality of the focal factors.

## Discussion

We found that fast explorers overall were more willing to consume bitter crickets and quickly stopped prioritizing on normal-tasting crickets in later learning trials, when there was no uncertainty regarding the number of each prey type and the association between prey colour and taste. Conversely, slow explorers were more aversive to bitter crickets and did not further adjust their foraging behaviours much in the learning phase. Even though the colour-taste association was also predictable during the reverse learning, the experience during this phase nevertheless decreased the overall reliability of colour as an indicator of prey quality, which individuals had previously used for foraging decision-making. Slow and fast explorers generally responded in similar manners to this new environmental uncertainty, while maintaining their differences in foraging behaviours. Our results offered both an intriguing contrast and a valuable addition to the literature concerning personality-cognition correlations.

### No evidence for a learning speed-flexibility trade-off

The common sun skinks were overall quick to associate prey colour with taste and immediately prioritized on higher quality prey. At the end of reverse learning, most individuals were also able to adjust their foraging priority and level of avoidance to values similar to those at the end of the previous learning phase, despite the switched colour-taste combinations (figures 1 and 2). Our results therefore did not show a trade-off between learning speed and flexibility at the among-individual level. This finding is in agreement with the pattern revealed by a recent meta-analysis [9]. We are aware that our finding did not entirely refute the possibility of such a trade-off, which can be masked when individuals being compared differ consistently in the amount of energy acquisition (reviewed in [5]). It is possible that we will see a negative correlation between learning speed and flexibility when each individual is faced with limited energy intake, which will be an intriguing future direction. Nevertheless, our data do suggest that learning speed and flexibility may not be under hard-wired constraints as previously hypothesized (reviewed in [8]). Instead, these two cognitive traits might be able to vary independently and respond separately to selection in the common sun skinks, at least when individuals are not energetically challenged.

### The correlation between exploration and the proactive-reactive learning styles

The fact that slow and fast explorers differed more in initial learning was consistent with the literature ([32–34], but see [12,35,36]). Importantly, our data revealed a correlation between exploration and learning style that was more nuanced than the predictions from the slow-fast syndromes, with both matching and mismatching aspects. Consistent with the slow-fast syndromes, fast explorers were indeed less choosy about prey quality. Even though low choosiness could result from less precise association between prey colour and quality (e.g., [37]), this result may not necessarily mean that fast explorers were making less accurate foraging choices. Foraging outcome often reflects a dynamic decision-making process, and consuming more low-quality prey could be beneficial both in terms of energy intake and information acquisition [38]. Since fast explorers move more per unit time, consuming more prey likely help satisfy their higher energetic needs, especially when the low-quality prey in our experiments were merely unpalatable. On the other hand, fast-exploring skinks exhibited a more reactive learning style, as they modified their foraging priorities more with the accumulation of information, which is opposite from previous predictions. Furthermore, the reactive learning style was characterized by marked flexibility without being slow in first associating colour with prey quality, another contrast to the conventional definition [7]. Even though slow explorers were more proactive, as they modified their foraging behaviour less as information accumulates, they were nonetheless also quick to prioritize on normal crickets, a demonstration of reactiveness.

Despite the fact that the exploration-cognition correlation as predicted by the fast-slow syndrome was not supported, we do not think this fact categorically refutes the fundamental concept linking personality and learning style. Rather, we believe that our results highlight the importance for considering the covariation and coevolution of personality and cognitive traits in an ecologically appropriate context, which can vary based on the foraging habits and life-history of the focal species. We posit that both the shared and distinct features between the “fast” and “slow” couplings between exploration and learning style might represent two strategies for dealing with rapidly changing environmental certainties; this hypothesis supplements existing theories, which focus on how slow and fast strategies might be favoured under different environmental conditions. Since the common sun skink is a widely ranging, generalist forager, even the slow-exploring individuals would likely encounter a diverse array of prey. As such, being able to quickly discern prey quality with reliable visual features (when available) would be favoured regardless of an individual’s exploration tendency. This might explain the quickly modified priority towards higher-quality prey observed in both slow and fast explorers in the early learning trials. For fast explorers, which tended to consume the majority of the crickets, it might not be useful to maintain any foraging priority once enough information has been collected regarding the number of each prey type and the meaning of colour as an indicator of prey taste. For slow explorers, which avoided bitter crickets more, maintaining a priority for normal crickets while still sampling crickets with the “low-quality” colour might be more beneficial. Interestingly, slow and fast explorers both modified their foraging behaviours similarly after initial confusion at the onset of reverse learning, despite reaching to different behavioural end points. This indicates that individuals, regardless of their exploration tendencies, were able to return to previously adopted foraging decisions under the new “meanings” of colours.

### Sex and colour cue as mediators of behavioural changes through learning

Our results also revealed sex to be an important determinant of foraging behaviour and learning styles, even though it did not play a mediating role in the personality-cognition correlation. Compared to females, male common sun skinks were less choosy and did not show a clear foraging priority during reverse learning, (figure 3). We note that this observed sexual difference was largely unrelated to exploration, as there was only weak evidence for sexual differences in exploration tendencies. This finding contrasts with a recent meta-analysis, which reported sex as a key mediator between personality and cognitive traits, even though the authors cautioned the generality of this pattern [9]. Nevertheless, our study was among the relatively few that explicitly tested the role of sex in personality-cognition correlation. The only significant mediator for personality-cognition coupling in the common sun skinks was prey colour. Even though the mediating role of colour has rarely been explored, it is well reported that colours differ in their efficacy as an avoidance learning cue (e.g., [39–44]). The most remarkable mediating effect of colour was observed in slow explorers that initially associated red and then yellow with bitter prey. These individuals did not learn to prioritize on red crickets that now tasted normal and still attacked and consumed a substantial number of bitter yellow crickets throughout the reverse learning phase (figure 1a, 2a). As a previous study found that red colour induced stronger avoidance than yellow colour in this species [10], it is likely that the efficacy of red as a warning colour had a longer carrying-over effect on slow explorers, which were more conservative foragers to begin with. The mediating effect of different colours on personality-cognition coupling has rarely been investigated but could be ecologically important, as foragers in nature will encounter a wide variety of colours that may or may not be reliable indicators of prey quality.

## Conclusion

In this study, we reported the correlation between exploration and learning style in a generalist lizard predator. Even though the nature of this correlation does not fully conform to predictions from the conventional fast-slow syndrome, the observed coupling between exploration and learning style makes ecological sense considering the environmental variability that the focal species might encounter. Our study also offered rare and valuable evidence (or lack thereof) for the role of sex and prey colour in mediating the personality-cognition correlation. Going forward, an important next step would be to obtain mechanistic evidence of the environment as the driver of the (co)evolution between personality and cognition traits. As data a broader range of taxa become available, a comparative study would be appropriate to test the causal relationship between environmental conditions and the form of personality-cognition coupling. Experimental evolution experiments that manipulate relevant environmental factors (e.g., food availability, reliability of colour as indicator for food quality, etc) would also yield valuable insights.

## Acknowledgements

We thank Hui-Chi Liu for her help with animal collection.

